# Intraspecific variability rivals interspecific differences in root traits of temperate tree seedlings

**DOI:** 10.64898/2025.12.15.694302

**Authors:** Morgane Dendoncker, Audrey Maheu, Monique Weemstra, Olivier Villemaire-Côté, Léa Darquié, Guillaume Lobet, Nelson Thiffault, Anne Ola, Alison D. Munson, Gabriel Asselin, Christian Messier, Isabelle Aubin

**Author notes:** **Corresponding author:**. Present address of the corresponding author: Earth and Life Institute, Université catholique de Louvain, Croix du Sud, 2 box L7.05.24, Louvain-la-Neuve, 1348, Belgium.

## Abstract

Global change and associated disturbances are increasing the risk of regeneration failure for tree species in temperate forests. Seedlings are particularly vulnerable to water stress due to their shallow root systems, making belowground plasticity a potentially key component of species adaptive capacity. Quantifying root trait variability and its drivers can improve our understanding of regeneration success under increasingly warm and dry conditions.

We quantified between species variation (BTV) and intraspecific variation (ITV) in seven root traits linked to water uptake—root-to-shoot ratio, maximum rooting depth, proportion of absorptive roots, specific root length, root tissue density, average absorptive root diameter, and root branching density—for seedlings of seven common, co-occurring tree species in forests of northeastern North America. We sampled seedlings under contrasting climate and light conditions, and assessed the influence of abiotic (climate, light conditions, soil properties) and biotic drivers (neighboring vegetation) as well as seedling characteristics (species identity, age, spermatophyte type) on root ITV at local and regional scales.

Species differed significantly for some traits but differed even more strongly in multivariate trait syndromes, suggesting distinct belowground strategies. ITV was substantial but trait-dependent, with maximum rooting depth and root to shoot ratio being the most variable (coefficient of variation > 45%) and branching density the least variable. BTV was the primary driver of overall trait variation for three traits, explaining more than 60% of variation, whereas within-plot ITV accounted for more than 50% of variation in the remaining four traits. Local drivers did not outweigh regional factors, and the overall explanatory power of measured drivers was limited, suggesting that fine-scale heterogeneity, not captured in our study, may strongly influence root ITV.

High ITV in most traits suggests substantial plasticity in roots, which may contribute to the adaptive capacity of seedlings facing climate change. Integrating this plasticity into mechanistic models is critical for predicting regeneration dynamics or root-mediated ecosystem processes. We propose a set of guidelines for integrating root traits into comparative studies and models based on trait measurability and extent of ITV. We further highlight the need to account for the scale- and gradient-intensity dependence of ITV-environment relationships.

## 1. Introduction

Anthropogenic global change brings an increased number of disturbances to forest ecosystems across the globe (McDowell *et al*., 2020; Seidl *et al*., 2017), including temperate forests (Sommerfeld *et al*., 2018). Different types of disturbances (e.g., insects, pests, drought, fires) can trigger shifts towards alternative vegetation structure and composition (Bowd *et al*., 2023; Millar and Stephenson, 2015). Cumulative or interacting disturbances may particularly disrupt forest dynamics by affecting regeneration success as seedlings, constrained by shallow root systems and limited energy reserves, are less able to withstand stress than mature trees (Niinemets, 2010; Walck *et al*., 2011). Under combined stressors such as herbivory (Beguin *et al*., 2016; Royo and Carson, 2022), fire (Boucher *et al*., 2020; Cerioni *et al*., 2024), and drought (Boucher *et al*., 2020; Crockett and Hurteau, 2024), regeneration failure poses an important threat to temperate forest equilibrium (Vickers *et al*., 2019).

In northern temperate forests, water stress is a major constraint for the establishment of both naturally regenerated and planted seedlings (Grossnickle, 2005; Walters *et al*., 2023). As the primary organs for water uptake, roots and their traits are closely linked to seedling survival ((Grossnickle, 2012). While root functional ecology has advanced, most research has focused on interspecific comparisons (Bergmann *et al*., 2020; Laughlin *et al*., 2021; Weigelt *et al*., 2021) with less attention to intraspecific trait variation (ITV). Yet, incorporating ITV is essential for understanding ecosystem dynamics and species responses to disturbances (Violle *et al*., 2012; Westerband *et al*., 2021), particularly under climate change (Aubin *et al*., 2016; Schaffer-Morrison *et al*., 2024). Few studies have examined root ITV in temperate forests (Comas and Eissenstat, 2009; Schaffer-Morrison *et al*., 2024; Tobner *et al*., 2013; Weemstra *et al*., 2017; Zadworny *et al*., 2016). A greater interspecific than intraspecific variation was found for root morphological and architectural traits by Comas and Eissenstat (2009), suggesting distinct species acquisition strategies. Other studies have reported that the degree of ITV differed between traits (Tobner *et al*., 2013; Weemstra *et al*., 2021; Zadworny *et al*., 2016), with biomass allocation showing plasticity across temperature and soil gradients, while root morphological traits were less responsive (Tobner *et al*., 2013; Weemstra *et al*., 2017). The direction and strength of ITV in roots also appears to be species-specific (Weemstra *et al*., 2021), highlighting the need for multispecies studies to identify the multiple ways through which plants can adjust their belowground strategies to different environmental conditions. Finally, organism-level traits such as root mass fraction and rooting depth – key for water and nutrient uptake – remain understudied though their plasticity is likely high according to the functional equilibrium theory which suggests that plants adjust biomass allocation to roots and shoots to balance acquisition of limiting resources (Brouwer, 1963; Poorter and Nagel, 2000).

Canopy openings induce substantial microclimatic alterations for the forest understory. By increasing light exposure at ground level, near-surface temperature is raised (Brackett *et al*., 2024), exposing seedlings to higher vapor pressure deficits, which can cause water stress. The role of light in shaping leaf trait plasticity is well established (Keenan and Niinemets, 2016; Niinemets *et al*., 2015; Poorter *et al*., 2009), but its influence on root traits – especially in trees – remains understudied (although see Poorter and Ryser, 2015). According to the functional equilibrium hypothesis (Brouwer, 1963), increasing light availability would decrease root-to-shoot biomass allocation. Indirectly, these changes modify competitive interactions for light and below ground resources among understory vegetation and co-occurring tree seedlings, with likely consequences for the root spatial distribution and traits of tree seedlings (Balandier *et al*., 2022). Understanding how light availability and understory affect root traits is crucial, especially for species that regenerate after canopy-altering disturbances or through reforestation efforts.

To improve our understanding of tree regeneration in increasingly dry and warm temperate forests (Huntington *et al*., 2009), our study aims to quantify the root ITV for seedlings of seven common, co-occurring tree species in the forests of northeastern North America, under different climate and light conditions representing the environmental gradient they typically occupy. Specifically, we addressed the following questions:

(Q1) How large is the variability within species (i.e., between- and within-plots) compared to between-species across root traits and root trait categories? We hypothesize that (H1a) species identity (i.e., between-species trait variation - BTV) is the dominant contributor to total trait variation, followed by ITV within plot and ITV between plots with species-specific and trait-specific differences in the variance structure; and (H1b) root system traits are more plastic than morphological and architectural traits.

(Q2) What are the drivers (climate, light availability, soil properties, neighboring vegetation and seedling characteristics) of root ITV across species, and at which spatial scale (individual-, plot-, region-level) do they operate? We hypothesize that (H2) local drivers (abiotic and biotic) are the dominant influences on ITV compared to regional drivers such as climate.

We tested the effects of (i) an abiotic regional driver (climate); (ii) abiotic local drivers (light conditions, soil properties); (iii) a biotic local driver (neighboring vegetation); and (iv) seedling characteristics (species identity, age, spermatophyte type) on both interspecific and intraspecific variation in seven root traits linked to water uptake. These root traits included three root system traits (root to shoot ratio, maximum rooting depth, proportion of absorptive roots; McCormack *et al*., 2017), three morphological root traits (specific root length, root tissue density, average diameter of absorptive roots), and one architectural root trait (root branching density). In comparison, one foliar trait (specific leaf area) was included to assess whether the relationships between the measured drivers and trait values remain similar between belowground and a key aboveground trait.

## 2. Material and methods

### 2.1. Replication statement

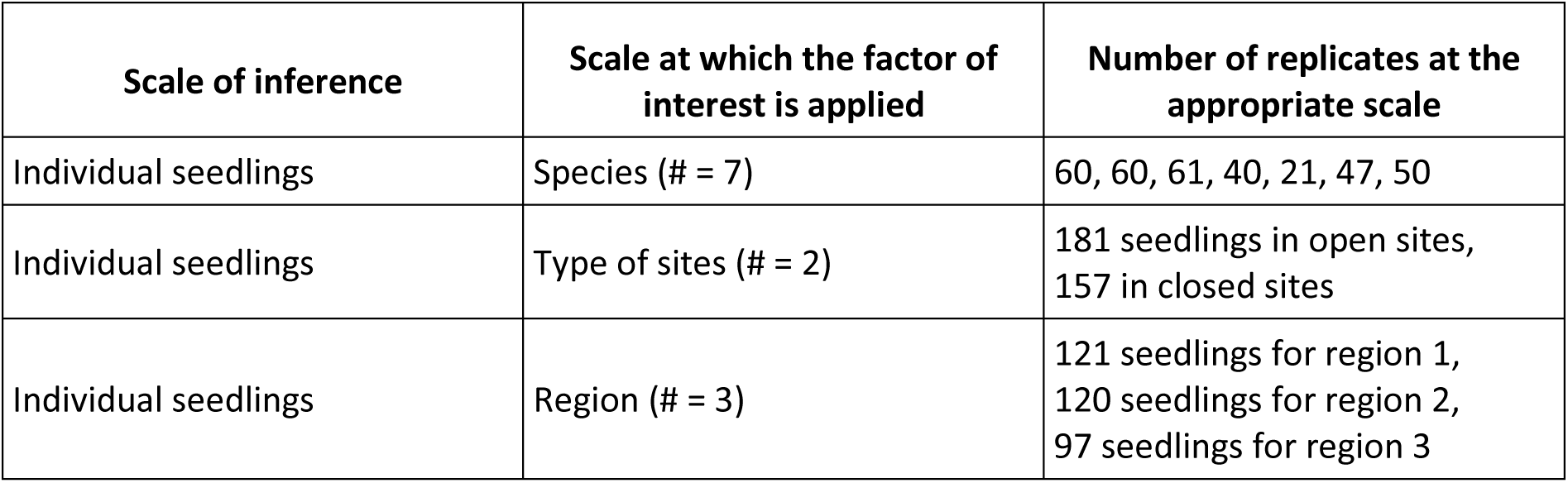

### 2.2. Study species and sampling design

We focused on seven abundant, co-occurring species of the temperate forests of northeastern North America, at the seedling stage: two gymnosperms (*Abies balsamea* (L.) Mill., *Pinus strobus* L.) and five angiosperms (*Acer saccharum* Marshall, *Acer rubrum* L., *Betula alleghaniensis* Britton, *Betula papyrifera* Marshall, *Quercus rubra* L.). Seedlings were defined as individuals that emerged directly from a seed (i.e., no resprouts nor root suckers), were upright and measured between 10 and 30 cm in height (from root collar to apex). We excluded creeping stems with adventitious roots and browsed seedlings identified by their thick stems with regrowth.

Seedlings were sampled using a hierarchical nested design (Figure 1). Fourteen plots, representing two contrasting light conditions (open vs. closed sites), were distributed across three Canadian regions with distinct climate and soil conditions (Supplementary material SM1): (1) Petawawa (Ontario; 46.00°N, 77.43°W); (2) Kenauk (Quebec; 45.75°N, 74.83°W); and (3) Duchesnay (Quebec; 46.60°N, 71.40°W). Mean annual precipitation (1984–2024; ERA5 land reanalysis; Hersbach *et al*., 2020) was 960 mm in Petawawa, 1100 mm in Kenauk, and 1350 mm in Duchesnay.

**Figure 1.**
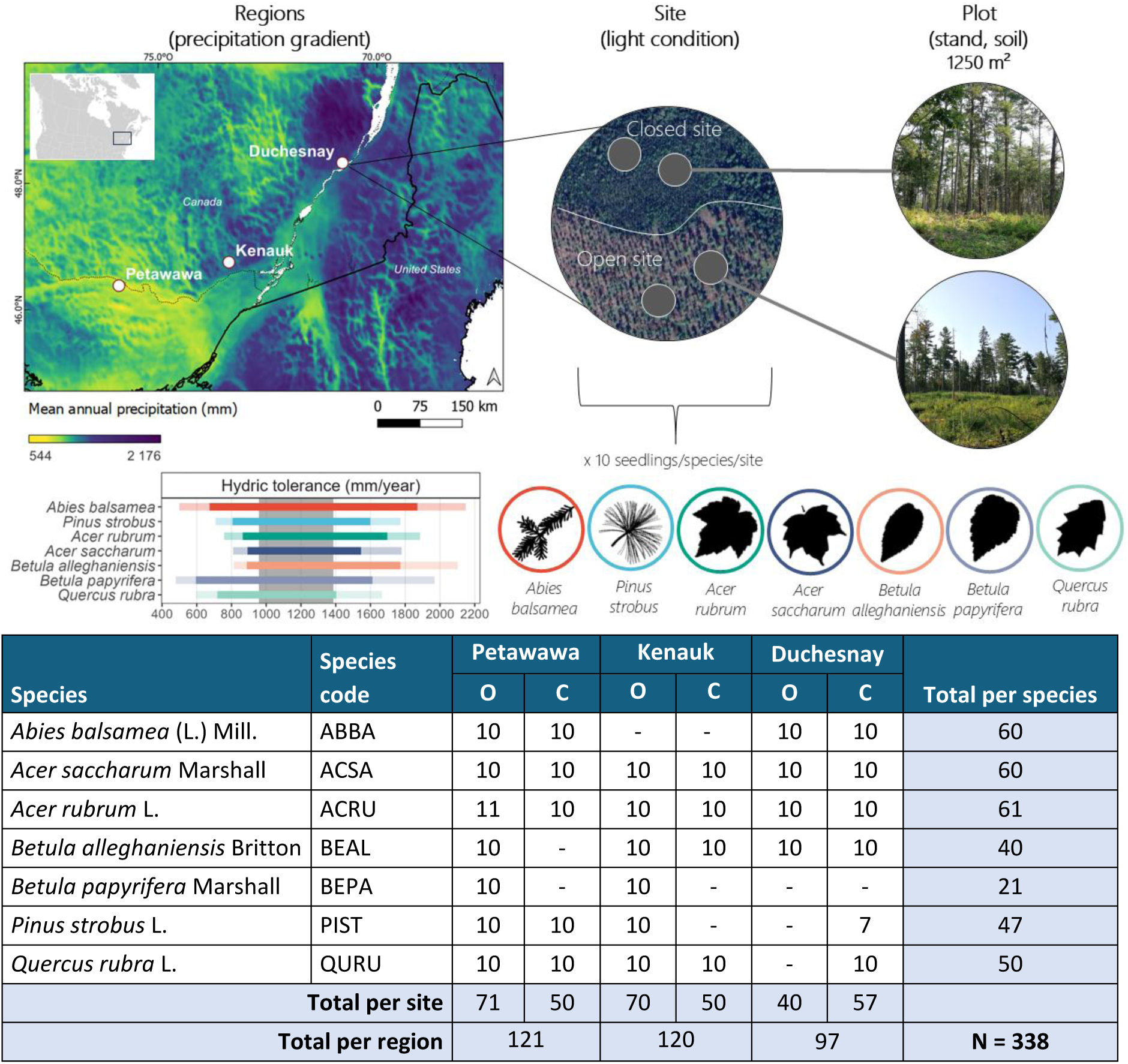
Sampling design. The seven species were sampled across 14 plots in three regions (Petawawa, Kenauk, Duchesnay), representing a precipitation gradient. Precipitation data was obtained from CHELSA dataset V2.1. (Karger et al., 2021; Karger et al., 2017). Each region included open vs. closed sites to reflect light conditions. Forest stand, understory, and soil features were measured at the plot scale. Species hydric tolerance (transparent bars: 1st–99th and solid bars: 5th–95th percentiles) is compared to the annual precipitation gradient covered by the three regions (grey rectangle). The table shows seedling counts per species, region, and light condition (O: open; C: closed).

Each plot (∼1250 m²) was flat or gently sloped (<5%), with similar species composition and vegetation structure. Plots were categorized by light condition: a closed site designated a mature forest with a closed canopy and no recent major disturbance; an open site referred to an opening of at least 30 m wide, resulting from a partial cut or natural disturbance. This threshold corresponds to the highest regeneration failure risk in temperate hardwood forests (Knapp *et al*., 2021) and reflects realistic conditions for partial cuts in northeastern North America. Gaps dominated by early successional small shrubs species such as *Rubus* spp. were excluded.

### 2.3. Data collection

Field work was conducted between July 29 and August 9, 2024. For each plot, forest stand structure was assessed within a 400 m² circular area. All trees with a diameter at breast height (1.3 m) greater than 5 cm were recorded to compute basal area (m².ha-1). Understory vegetation cover was surveyed using four 1 m² quadrats, positioned 5 m from the plot center at the four cardinal points. Taxa covering more than 5% of the quadrat area were identified to the genus (forbs) or family/order (ferns). Humus depth was measured near the plot center. One mineral soil sample per plot was collected at the same location at a 20 cm depth for texture analysis and total carbon (C) and nitrogen (N) concentration using a Thermo Scientific Flash 2000 Elemental Analyzer.

We sampled 10 seedlings per species for each light condition–region combination, following the selection criteria described above by sweeping the plots outwards from the center. There were a few exceptions to this number: *Betula* spp. which were rare in closed sites, and *Pinus strobus* and *Quercus rubra*, which were absent from open sites in Duchesnay (Table 1). Each seedling root system was fully excavated. The local microtopography was characterized and recorded for each seedling (Liechty *et al*., 1997), where pits were features with a depth of at least 40 cm (bottom below the general soil surface); mounds were at least 40 cm above the general soil surface and consisted primarily of mineral soil; and flat were all other features not meeting the criteria of a pit or mound.

**Table 1.** Investigated drivers of root trait variability with their associated variables and observed range (or levels for categorical variables). Drivers are grouped by type (abiotic regional, abiotic local, biotic local and individual), which reflects their nature and spatial scale.^1^

| Type | Driver | Scale | Variable | Range/levels |
| --- | --- | --- | --- | --- |
| Abiotic regional | Climate | Region | CMI: average annual P-PET over 2020-2024 (mean over 2020-2024) | 360 – 790 mm |
|  |  |  | Average annual air temperature (mean over 2020-2024) | 6.1 – 6.9°C |
|  |  |  | May-September P-PET (mean over seedling life span) | 350 – 850 mm |
|  |  |  | Average annual air temperature<br>(mean over seedling life span) | 5.3 – 8.2°C |
| **Abiotic local** | Microtopography | Individual | Microtopography | Pit, mound, flat |
|  | Light condition | Plot | Canopy openness | Open, closed sites |
|  | Soil properties | Plot | Soil texture | Sand: 52 – 97%<br>Silt: 1 – 38%<br>Clay: 1 – 17% |
|  |  |  | Humus layer depth | 0 – 13 cm |
|  |  |  | Total C concentration | 0.3 – 9.2% |
|  |  |  | Total N concentration | 0.01 – 0.5% |
| **Biotic local** | Understory | Plot<br>(average<br>over 4<br>quadrats) | Proportion of ferns, forbs,<br>graminoids | Ferns: 0 – 100%<br>Forbs: 0 – 100%<br>Graminoids: 0 – 71% |
|  |  |  | Percent cover | 30 – 100% |
|  | Forest stand | Plot | Basal area | 0 – 55.6 m<sup>2</sup>/ha |
|  |  |  | Proportion (in basal area) of<br>angiosperms trees (%) | 37 – 100% |
| **Seedling**<br>**characteristics** | Age | Individual | Seedling age | 1 – 18 years |
|  | Phylogeny |  | Seedling spermatophyte clade | Angiosperm,<br>Gymnosperm |

<sup>1</sup> CMI: Climate Moisture Index; AP: annual precipitation; PET: potential evapotranspiration; C: carbon; N: nitrogen; P, precipitation

After root excavation, the following measurements were taken for each seedling: (i) total height (i.e., length of the main stem, from root collar to apical bud); (ii) the crown width in two perpendicular directions; (iii) the root collar stem diameter; (iv) the maximum rooting depth (i.e., the depth at which the deepest root ends); and (v) the maximum root lateral spread (i.e., the maximum horizontal distance at which a lateral root is found). Seedling age was estimated by counting terminal bud scars for angiosperms, and tree rings under a binocular microscope for gymnosperms.

The root system was then separated from aboveground parts. For gymnosperms, needles and stem were kept together; for angiosperms, leaves were separated from the stem. Each compartment (roots, stems, leaves) was wrapped in moist paper, placed in a plastic bag and stored in an insulated cooler until transported to the lab at the end of each day. Samples were refrigerated at 4°C up to one week before further processing.

### 2.4. Trait measurements

To determine leaf area, angiosperm leaves were scanned with a flatbed scanner at 300 dpi. For gymnosperms, needles were separated from stems, and 20 randomly selected needles were scanned at the same resolution. All leaves and needles were oven-dried at 60°C for 48 h. Stems were dried at 60°C for 72 h. Dried leaf and stem tissues were weighed using a Pioneer analytical balance (Ohaus PX323, precision ± 0.001 g). Specific leaf area (SLA; cm².g^-1^) was calculated for (i) angiosperms as total abaxial leaf area (excluding petioles) divided by dry mass and (ii) gymnosperms as total area of the 20 scanned needles divided by their dry mass.

Roots were washed to remove soil particles and stored in 50% ethanol at 4°C until scanning (Freschet *et al*., 2021). Root systems were scanned at 1200 dpi on an EPSON Expression 11000XL scanner, providing a resolution of 47 pixels.mm-1—suitable for fine root trait analysis such as root diameter. Two scan types were performed per seedling. The first included the whole root system (i.e., all roots including both fine and coarse roots). The second type focused on 10 fragments of absorptive roots, defined as first- and second-order roots, following morphometric classification where the first order corresponds to the most distal order (Freschet *et al*., 2021; Freschet and Roumet, 2017; McCormack *et al*., 2015). All scanned roots were oven-dried at 60°C for 72 h. Whole root systems were weighed on the Pioneer scale; absorptive root fragments were weighed on a Sartorius Cubis scale (Model MSA3.6P-0TR-DM, precision ± 0.001 mg).

Root scans were processed in ImageJ to remove edges and soil particles and enhance contrast. Binary images were analyzed in RhizoVision Explorer (Seethepalli *et al*., 2021; Seethepalli and York, 2020) to extract architectural and morphological traits. Following general root trait classifications (McCormack *et al*., 2017), we computed three root system traits. Root:shoot ratio (R:S; unitless) was calculated as the dry mass of the complete root system divided by the dry mass of the aboveground parts. The proportion of absorptive roots (PAR; %) was calculated by dividing the total length of roots below a certain diameter threshold (angiosperms: 0.5 mm; gymnosperms: 1 mm; based on the average diameter of the ten absorptive root fragments) by the total root system length. Maximum rooting depth (MRD; cm) was measured in the field as the soil depth reached by the deepest root. Three morphological root traits were computed from the absorptive root fragments. Root tissue density (RTD; g.cm^-3^) corresponds to the dry mass divided by root volume (from RhizoVision). Specific root length (SRL; m.g^-1^) is the total root length divided by dry mass. Average diameter of absorptive roots (AD; mm) was measured in RhizoVision. Finally, root branching density (RBD; tips.cm^-1^) – an architectural root trait – was calculated on the complete root system as the number of root tips divided by total root length (RhizoVision). These traits were selected for their relevance to woody seedling responses to water deficit (Supplementary material SM2).

### 2.5. Statistical analysis

To quantify the extent of intraspecific trait variation relative to between-species variation (Q1), we applied both single-trait and multi-trait approaches. For the single trait approach, ITV was first assessed by calculating the coefficient of variation (CV) per trait, per species, which was then compared across traits with an ANOVA and Tukey’s HSD post-hoc test. Second, to compare species mean root trait values, we fitted linear mixed models (LMMs) using the lmerTest R package (Kuznetsova *et al*., 2017; R core team, 2022) with species as a fixed effect and region and plot nested as random effects. We then calculated estimated marginal means for each species-trait combination using the R package emmeans (Lenth, 2023) and performed post-hoc Tukey’s HSD tests to assess pairwise differences.

The multi-trait approach was applied to determine whether species can be differentiated based on their root trait syndrome (i.e., the combination of trait values), using a Linear Discriminant Analysis (LDA). This analysis identifies linear combinations of continuous variables (here, root traits) that best separate groups (here: species; Legendre and Legendre, 2012), allowing us to estimate the probability of correctly assigning seedlings to their respective species based on trait profiles. A high probability of correct assignment would support hypothesis H1a.

Finally, to assess the relative contribution of species identity, and within- and between-plot variation to overall trait variation (H1a), we performed variance partitioning using LMMs. For each trait, we fitted models with only random factors: (i) species (i.e., between-species variation) nested in (ii) plots (i.e., ITV between plots) nested in (iii) regions (i.e., ITV between regions). Additionally, we fitted separate models for each species and each trait, to assess potential species-specific differences in variance structure. The two *Betula* spp. were excluded from the latter analysis due to their uneven distribution across all regions and light condition combinations.

To examine the influence of biotic and abiotic drivers at both local and regional scales, as well as seedling characteristics, on ITV (Q2), we fitted single-trait LMMs. Random intercepts included region, plot nested within region, and species, to account for species-level variation and focus on intra-specific trait variation. Fixed effects encompassed abiotic regional variables (climate); biotic local (neighboring vegetation); abiotic local (microtopography, light conditions and soil properties) and seedling characteristics (age and spermatophyte clade, Table 1). Climatic variables were derived from ERA5-land reanalysis data (Hersbach et al., 2020). For each region, we calculated the Climate Moisture Index (annual precipitation, P, minus potential evapotranspiration, PET) and the average annual air temperature over the period 2020-2024 to characterize regional climate. These years correspond to the overall age of the seedlings considered in the study. Additionally, we computed climate conditions of the growing season for each seedling, over the period corresponding to its age, to reflect the actual climate conditions experienced by each individual.

For each trait, an initial complete model was constructed including all fixed effects. Variance Inflation Factors (VIFs) were calculated to assess multicollinearity, and variables with high VIFs (> 5) were excluded. Model selection was then performed using the step function of the lmerTest package in R (Kuznetsova *et al*., 2017), which applies a backward elimination of random and fixed effects and compares models via ANOVA. Assumptions of homoscedasticity, independence and normality of residuals were visually inspected; some traits were log-transformed to meet these assumptions. Pseudo R² values – marginal (R²m) and conditional (R²c; Nakagawa and Schielzeth, 2013) – were estimated using the r.quaredGLMM function from the MuMin package (Barton, 2020). They respectively represent the variance explained by fixed effects alone (R²m) and by both fixed and random effects combined (R²c). Semi-partial marginal R² values, indicating the contribution of each fixed effect (although fixed effects can share variance; Jaeger *et al*., 2017), were computed using the r2beta function from the r2glmm package (Jaeger, 2025). All continuous variables were scaled and centered prior to analyses.

## 3. Results

### 3.1. Between species trait variation

Trait means and distributions differed significantly across species (Figure 2; Supplementary material SM3). For example, the distributions of R:S were different across species and generally presented different means (Supplementary material SM4). *Quercus rubra* had a significantly higher R:S (1.1) than the two *Acer* species (0.70 and 0.80 for *Acer rubrum* and *Acer saccharum*, respectively). *Abies balsamea* and the two *Betula* species had similar, intermediate R:S (0.34-0.42), and *Pinus strobus* the lowest (0.26). Similarly, the distributions and means of RBD, SLA and SRL also differed widely, while MRD and PAR were largely similar across species.

**Figure 2.**
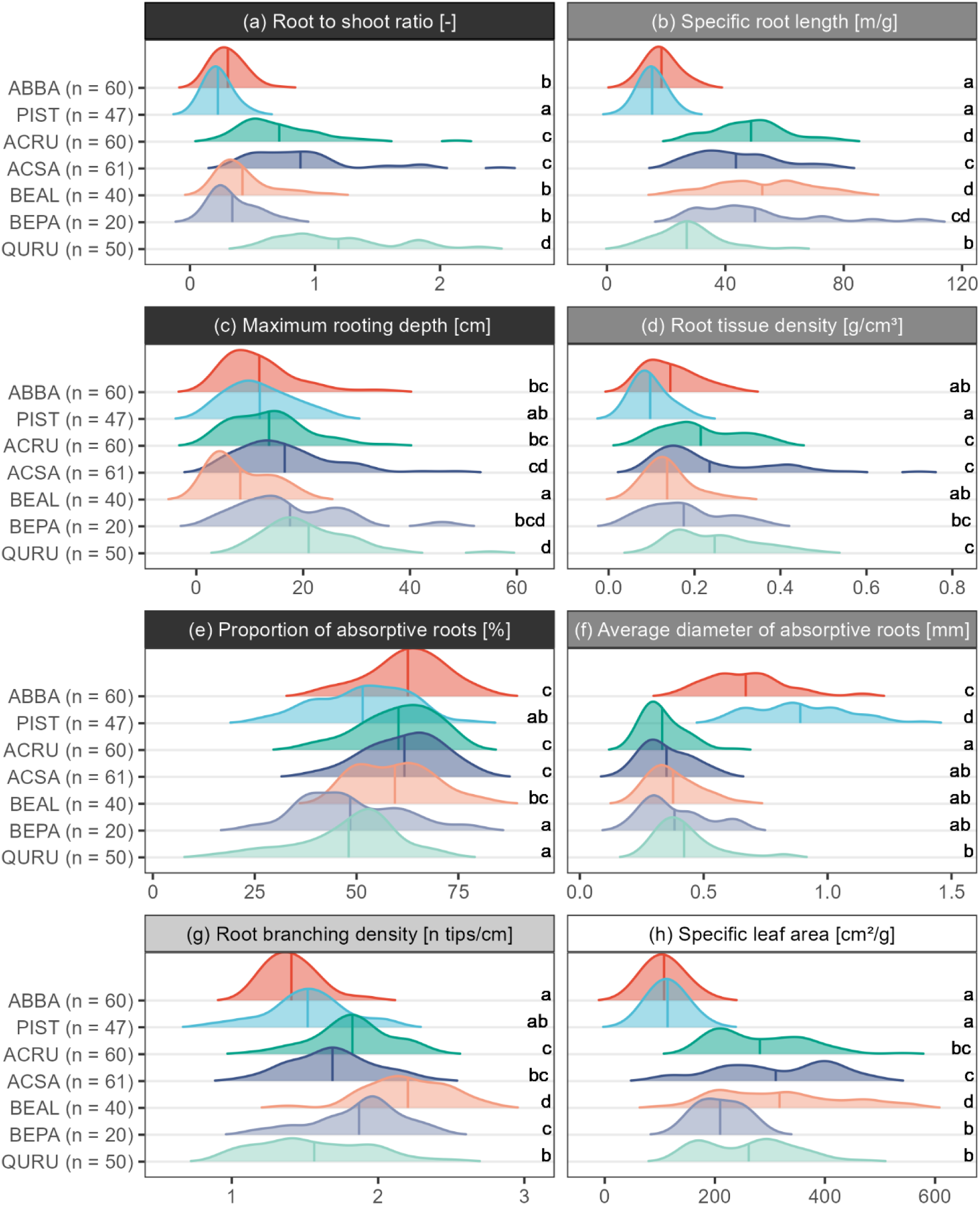
Estimates of density of seedlings’ traits per species (number of individuals between brackets, see Figure 1 for species abbreviations). The background color of the facet’s titles shows the type of traits: black = root system traits; dark grey = morphological traits; light grey = architectural trait; white = aboveground trait. Vertical lines represent the means of trait distribution. Means with different letters differ significantly (α = 0.05).

Across all traits combined, the first linear discriminant was mostly associated with the inverse of AD and, to a lesser extent, with RTD, and weakly with RBD (Figure 3a). This axis essentially separated the gymnosperms from the angiosperms. The second linear discriminant was explained by AD, R:S and RBD. The LDA correctly identified seedlings to the species level based on their root trait syndromes in 67.2% of the cases (Figure 3). Species with the highest classification accuracy were *Abies balsamea* (83.3%), *Quercus rubra* (78.0%), and *Betula alleghaniensis* (72.5%, Figure 3b). *Pinus strobus*, *Acer saccharum*, *Betula papyrifera* and *Acer rubrum* had a lower classification accuracy of 63.8%, 59.0%, 55.0% and 53.3%, respectively. Overall, gymnosperms were never misclassified as angiosperms (except for one individual of *Abies balsamea* misclassified as *Acer saccharum*) and vice-versa. Within genera, most of the misclassified *Acer rubrum* individuals were wrongly identified as *Acer saccharum* (16 individuals); 17 of 61 *Acer saccharum* individuals were incorrectly classified as *Acer rubrum*; and 5 of 20 *Betula papyrifera* individuals were wrongly classified as *Betula alleghaniensis*.

**Figure 3.**
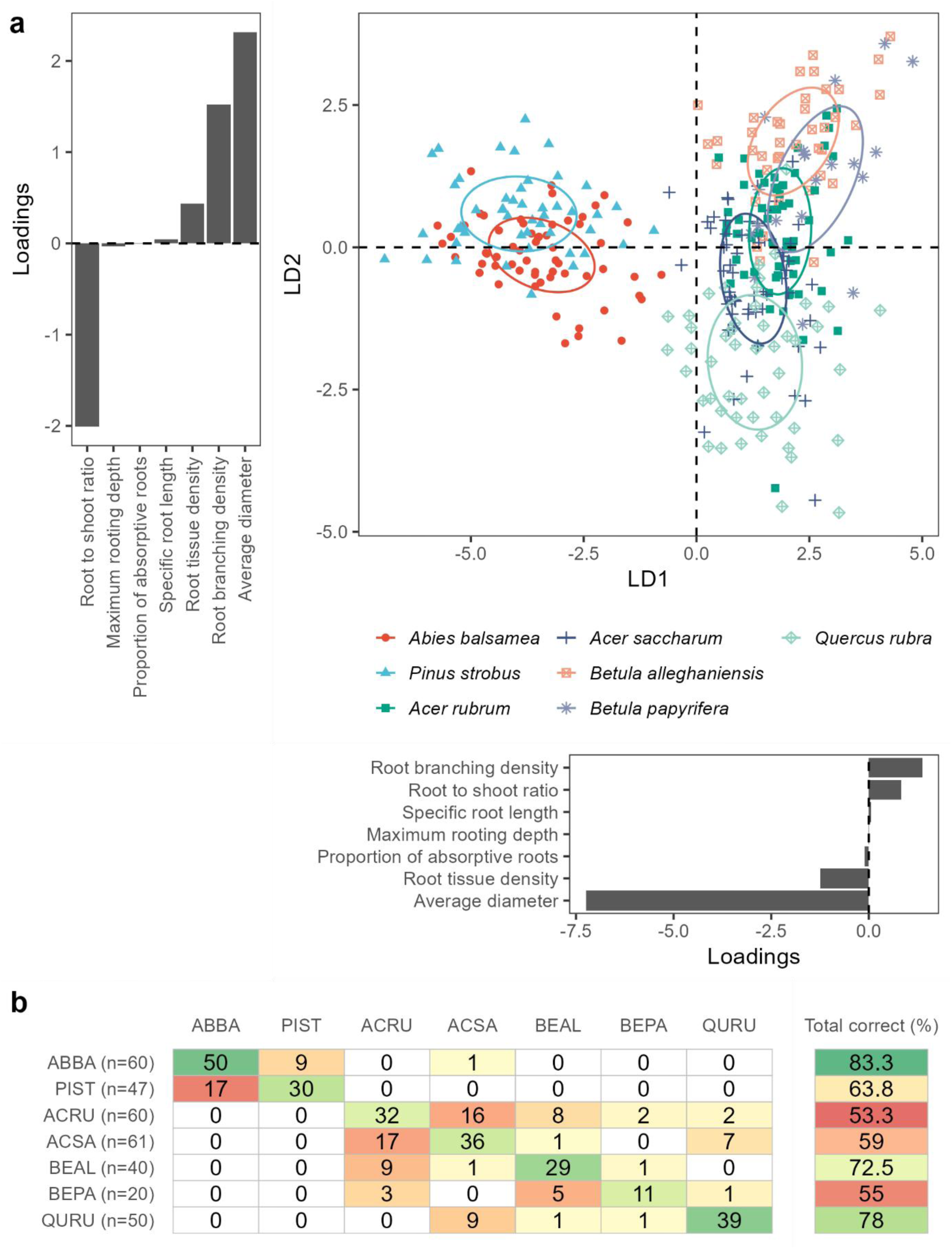
Linear Discriminant Analysis on root traits. (a) Seedlings (colored by species) in the first two dimensions (linear discriminants, LD) with associated loadings of the traits for LD1 and LD2. Ellipses represent 95% confidence regions for each species. (b) Contingency table with the observed species in rows and predicted species from the LDA in columns (see Figure 1 for species abbreviations). The final column indicates the percentage of individuals that were correctly classified; the darker the green, the higher the correct classification and the darker the red, the lower the correct classification.

### 3.2. Intraspecific trait variation

Overall CVs (i.e., averaged per trait across all seven species) differed significantly (Figure 4; black dots). MRD and R:S had the highest mean CV (54.1% and 45.7%), followed by RTD (44.1%), SRL (31.4%), SLA (29.3%), AD (26.7%), PAR (19.1%) and RBD (16.4%). Trait CV within species (Figure 4, solid bars) followed a similar order with highest CVs for MRD (from 40.0% for *Quercus rubra* to 65.2% for *Betula alleghaniensis*) and R:S (38.8% for *Abies balsamea* to 51.7% for *Betula papyrifera*), followed by RTD, SRL, AD. Lower variations were exhibited in RBD and PAR, with species’ CV always lower than 25%.

**Figure 4.**
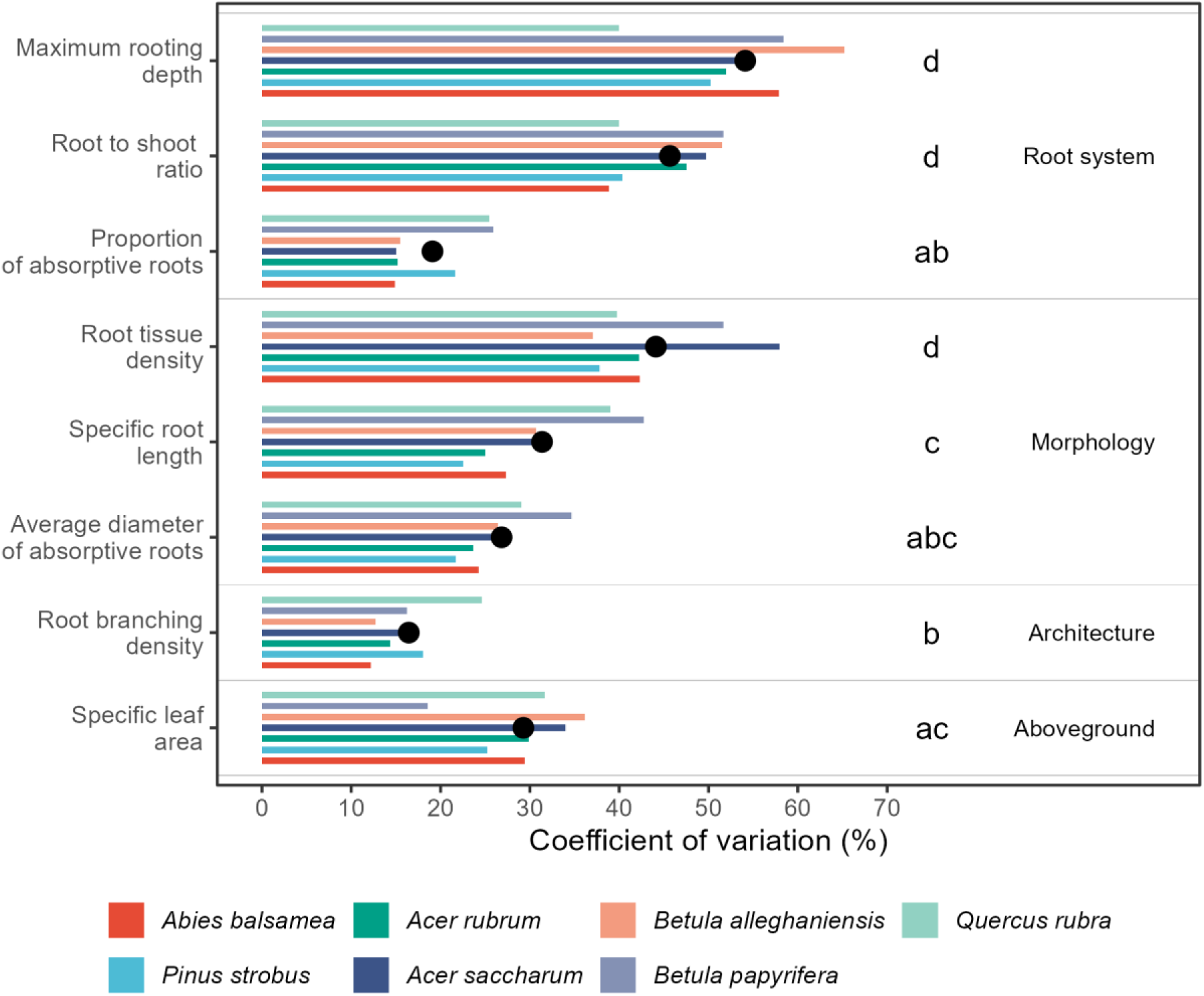
Trait coefficient of variation (CV) per species (solid bars) and overall mean CV across species (black dots). Different letters indicate significant differences in mean trait CV across species.

### 3.3. Variance structure of trait variation

The overall variation in traits (i.e., across all species) was mostly explained by two components: between species variation (BTV) and ITV within plot (ITVWP), i.e., root trait differences between individuals of the same species in the same plot; Figure 5a). BTV explained more than 60% of the overall trait variation for SRL (70.5%), AD (67.3%), R:S (64.4%) and SLA (61.4%). Overall variation in RBD was explained by ITVP (46.6%) and by BTV (38.1%); PAR by ITVP (53.3%) and by BTV (33.4%); and RTD by ITVP (66%) and by BTV (25%). BTV explained only a small proportion of the variation in MRD, for which ITVP was the main variance component (48.1%). ITV between regions (ITVBR, i.e., root trait differences between individuals of the same species across regions) generally explained less than 15% of the variation, except for MRD (33.8%). ITV between plots (ITVBP, i.e., root trait differences between individuals of the same species across plots) was also low for every root trait (less than 15%) but represented 26.8% of total variance for the SLA.

**Figure 5.**
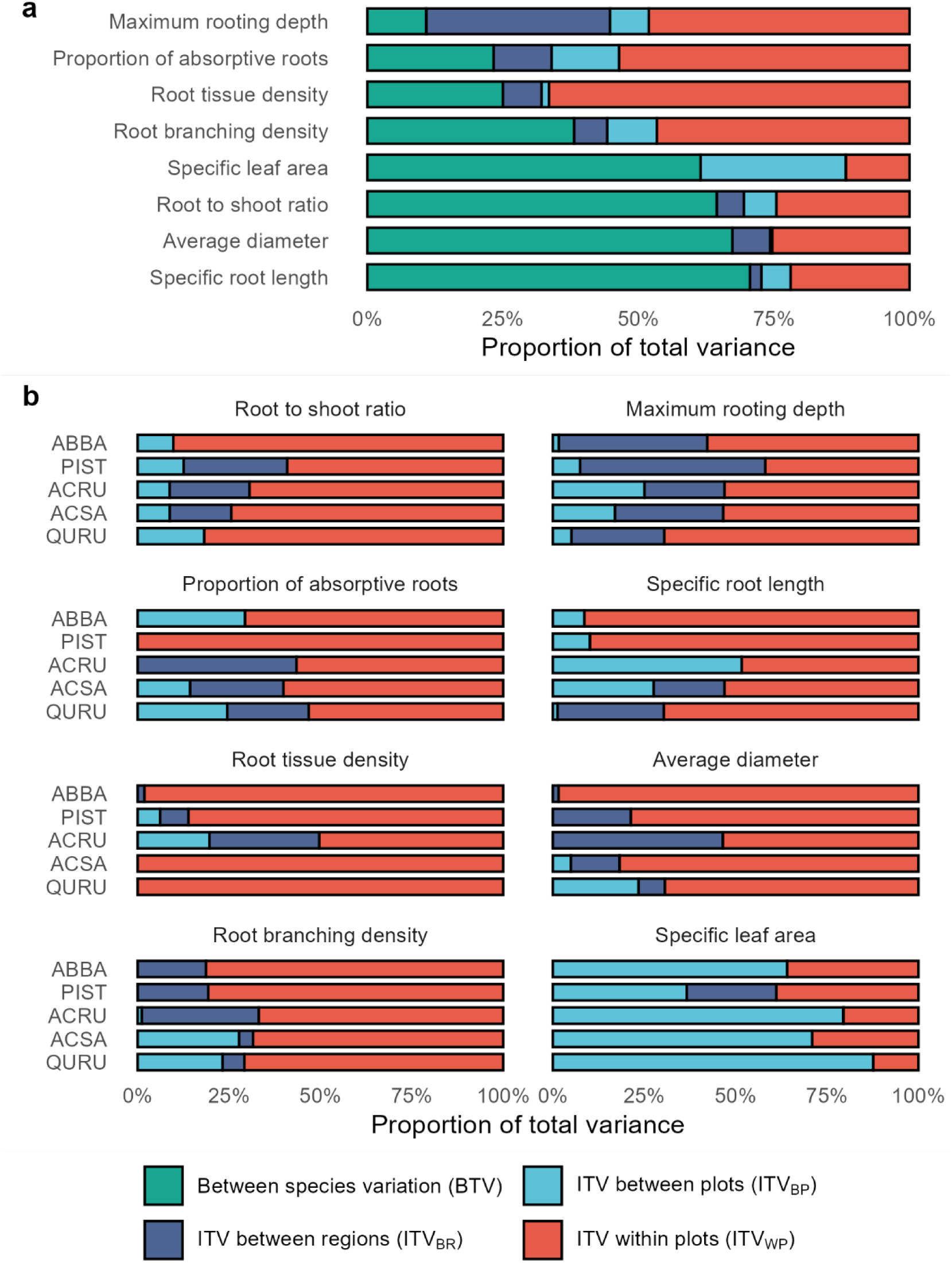
Proportion of total trait variance explained by species identity (between species variations, BTV), intraspecific trait variation between regions (ITV_BR_), intraspecific trait variation between plots (ITV_BP_) and intraspecific trait variation within plots (ITV_WP_). (a) Per trait, all species considered. (b) Per species, for Abies balsamea (ABBA), Acer rubrum (ACRU), Acer saccharum (ACSA), Pinus strobus (PIST) and Quercus rubra (QURU).

At the individual species level, different components structured the variance of different traits (Figure 5b). For all species and all root traits, more than 40% of trait variation was explained by ITVP, but the proportions explained by ITVBR and ITVBP differed. For instance, ITVBR explained more than 15% of R:S variance for *Acer rubrum*, *A. saccharum*, and *Pinus strobus*, but none for *Abies balsamea* and *Quercus rubra*. MRD had more than 20% of its variance explained by ITVBR across species. Unlike root traits, SLA had more than 60% of its variation explained by the ITVBP for all species, except for *Pinus strobus*. This species was the only one for which ITVBR explained a notable proportion of SLA variance (24.5%).

### 3.4. Biotic and abiotic drivers of root ITV

Intraspecific variation in individual traits was explained by different drivers (abiotic regional, abiotic local, biotic local and seedling characteristics; Table 1), and to variable extents, with marginal R² ranging from 0.01 for R:S to 0.79 for SLA (Figure 6a; Supplementary material SM5). Overall, with random effects considered, more variance was explained, with conditional R² ranging from 0.42 for PAR to 0.87 for SLA.

**Figure 6.**
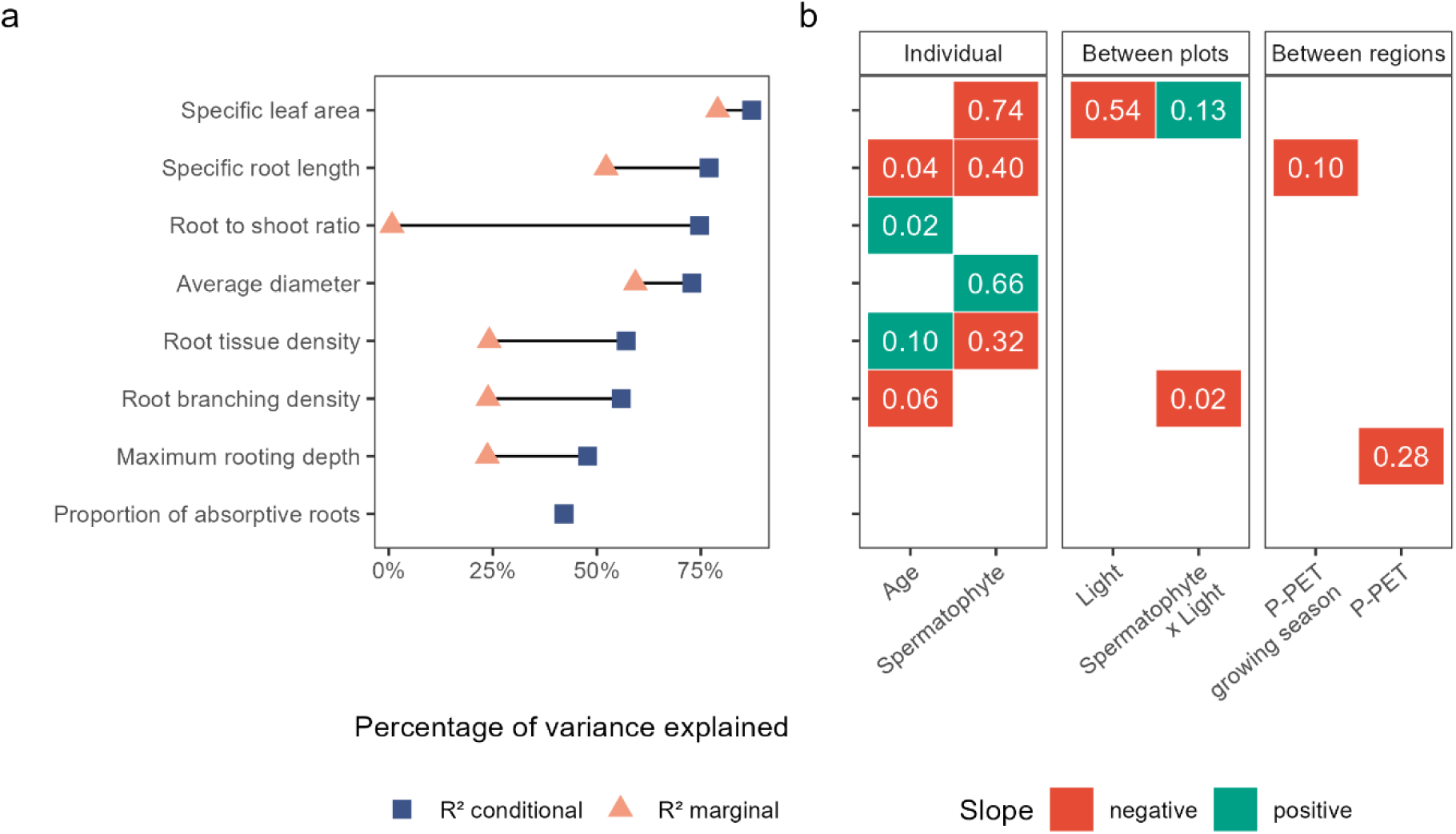
Drivers of ITV in root and leaf traits, and (a) the percentage of variance they explain (conditional and marginal R²) and (b) effect sizes of significant drivers semi-partial marginal R²m). Note that as fixed effects can share variance, the sum of all semi-partial R² values does not necessarily equal R²m. Red fill indicates a negative slope; green fill indicates a positive slope. For spermatophyte effects, negative/positive slopes indicate lower/higher values for Gymnosperms; for light conditions, negative/positive slopes indicate lower/higher values for open sites.

Fixed effects that explained the largest proportion of ITV were *seedling characteristics* (seedling age and spermatophyte clade, Figure 6b), significantly affecting six of eight traits. *Drivers at the plot and region scale*s (climate, soil properties, microtopography, light conditions, neighboring vegetation, Table 1) did not explain much of the ITV in all root traits, except for MRD. The ITV of this trait was driven by the difference in water balance (P-PET) across regions (semi-partial R²_m_= 0.28). ITV in SRL was also negatively associated with water balance across regions, but only weakly (semi-partial R²_m_ = 0.1). Finally, SLA variation was explained by spermatophyte clade, light conditions and their interaction.

## 4. Discussion

### 4.1. Between species trait variation rivals intraspecific trait variation

Global change and associated disturbances are increasing the risk of regeneration failure for some species in temperate forests, which may lead to shifts in forest composition and structure (Bowd *et al*., 2023). Seedlings are particularly vulnerable to water stress due to their relatively shallow root system (Tiebel *et al*., 2023). Importantly, tree species may differ, not only in their mean root trait values but also in the variability of these traits. In this study, conducted in the forests of northeastern North America, we found substantial between species trait variation (BTV) and intra-specific variation (ITV) in the root traits of seedlings from seven common temperate tree species. This ITV may confer adaptive advantages for some species under changing environmental conditions.

Firstly, consistent with previous studies, species differed significantly in their root traits. Supporting H1a, species identity emerged as the primary determinant of overall variation in root to shoot ratio (R:S), average diameter (AD) and specific root length (SRL), compared to ITV within and between plots and ITV across regions. However, root tissue density (RTD) and root branching density (RBD), maximum rooting depth (MRD) and proportion of absorptive roots (PAR) were predominantly influenced by within-plot ITV. The strong influence of species identity on total root trait variation of woody species has been documented in temperate forests for SRL, RTD and AD (Comas and Eissenstat, 2009; Valverde-Barrantes *et al*., 2013). Similarly, in their study on herbaceous and woody species along an elevation gradient Weemstra *et al*. (2021) found species identity to be the main source of variation for SRL, AD – consistent with our findings – and RBD – in contrast to our results. In line with this, LDA often correctly classified species based on their root trait syndromes (Figure 3). This suggests that co-existing species adopt distinct resource acquisition strategies.

At the same time, interspecific variation was – for most traits – not high enough to distinguish all species based on individual trait means (Figure 2). Thus, while species identity was a key driver of overall root trait variation, high ITV was also observed, although its extent varied across traits and species. For example, partly in line with Hypothesis H1b (predicting greater plasticity in root system traits compared to morphological and architectural traits): MRD and R:S, two of the three root system traits examined, exhibited higher ITV than two of the three morphological traits (SRL and AD). Previous studies have suggested greater plasticity in root system traits than in root morphological traits. For instance, Freschet *et al*. (2015) demonstrated a stronger adjustment capacity in herbaceous species to nutrient availability via changes in root mass fractions (root system trait, linked to R:S) rather than by their average SRL (morphological trait). Weemstra *et al*. (2017) also found higher plasticity in fine root biomass (root system trait) and lower plasticity in SRL and RTD (morphological traits) in beech and spruce forests established on contrasting soils. Contrary to our hypothesis, however, PAR (root system trait), exhibited the lowest ITV among the studied traits, while RTD (a morphological trait) showed plasticity levels comparable to MRD and R:S. The high ITV observed for R:S and the low ITV for PAR suggest that seedlings maintained a relatively stable pool of absorptive roots to ensure consistent water and nutrient uptake, while adjusting their biomass allocation in response to local environmental conditions.

Among the different drivers tested, several seedling characteristics influenced ITV (Table 1, Figure 6). For instance, the gymnosperm species had thicker roots, and lower SRL than the angiosperms. Since only two gymnosperm species were included in our sample, it would be premature to generalize this finding across all species in the studied regions, but these results agree with other studies that compared root traits between conifer and broadleaved tree species (e.g. Liese *et al*., 2017).

Age also had a significant, albeit small, effect on SRL, RBD, R:S and RTD. Ontogenetic and age-related effects on traits are well known (Barton, 2024; McCormack *et al*., 2017), even within ontogenetic stages, and particularly during the prolonged juvenile phase of forest trees. While our study did not specifically examine ontogenetic effects on root traits per se, we explored this aspect indirectly by comparing trait rankings between our seedling data and data from mature trees from the TRY (Kattge *et al*., 2020; Kattge *et al*., 2011) and TOPIC (Aubin *et al*., 2020) databases. While root mass fraction ranking – computed as proxy for R:S, for which data were unavailable for mature trees – stays relatively similar (Supplementary material SM6), changes in ranking were found for AD, SRL and RTD (e.g., *Pinus strobus* had the lowest mean RTD at seedling stage but the highest at mature stage). These ontogenetic root trait patterns may reflect a shift in water-use strategies as trees grow bigger (Cavender-Bares and Bazzaz, 2000; Di Iorio *et al*., 2024).

Finally, species exhibiting high plasticity for one trait did not necessarily show high ITV across all traits (Figure 4). For instance, *Quercus rubra*, which develops a taproot system, showed low plasticity for MRD, while this trait was highly plastic in the two *Betula* species. Conversely, *Quercus rubra* displayed high ITV in its root architecture (RBD) and morphology (SRL). This supports the ITV patterns reported across species in previous studies (e.g., Kumordzi *et al*., 2019; Weemstra *et al*., 2021), which further showed that root trait plasticity was far more idiosyncratic than that of leaf traits, and highlights the existence of species-specific belowground strategies.

### 4.2. Scale and gradient intensity dependence of root ITV drivers

Looking at the different drivers of ITV, we found no conclusive support for H2, which stated that local drivers would dominate regional abiotic effects on root ITV. Neither soil properties (abiotic local) nor understory vegetation and forest stand characteristics (biotic local) significantly affected ITV for any trait. This contrasts with previous studies which suggested that environmental influences on root traits operate at different spatial scales (Weemstra *et al*., 2021). Cardou *et al*. (2022) also documented significant effects of local drivers on the ITV of understory species traits in similar ecosystems across Canada, covering a much larger environmental gradient. Light conditions (abiotic local), however, did influence SLA and, to a lesser extent, RBD, with lower SLA and higher RBD observed under high light conditions. The interactions between light conditions, understory cover and composition might have mitigated their individual effects on root traits, as higher light promotes denser understory and subsequently increases competition for resources (Balandier *et al*., 2022). It is also important to note that light conditions were classified dichotomously as “open” versus “closed” sites, but there was a gradient of light availability within both classes. An experimental design that strictly compared full-light to near-complete absence of light might have produced a more pronounced contrasting response in root traits.

At the regional scale, water supply affected SRL and MRD, which both decreased under higher water availability, although its effect was limited (low pseudo-marginal R²). This finding is in line with results of Schaffer-Morrison *et al*. (2024), who also found small abiotic effects on the root traits of temperate tree seedlings, although in their work, local environmental effects exceeded regional climatic influences. The relatively small effect of water availability observed in this study is likely due to its relatively narrow gradient, especially when compared to more controlled and pronounced water treatments studied elsewhere (Jaeger *et al*., 2023; Jaeger *et al*., 2025; Olmo *et al*., 2014). Moreover, our study captured regional climate variations within a relatively homogeneous bioclimatic region in which all seven species co-occur. As a result, the precipitation gradient represented here spans distinct portions of each species’ climatic niche (Figure 1). This can obscure trait-environment relationships, as the climatic conditions in each region do not necessarily represent the optimal or limiting conditions for every species (Anderegg, 2023). For species with broader hydric niches, such as *Abies balsamea* and *Betula papyrifera* (Figure 1), it is likely that including a wider precipitation gradient would have revealed different trait responses.

Despite the limited influence of local drivers on the studied traits, we cannot conclude that regional drivers exert a stronger influence on the root ITV of the seven study species. Given the high proportion of variance explained by within-plot ITV – compared with ITV across plots and across regions – it is more likely that the relevant local drivers were not captured in our analysis. For example, soil phosphorous availability was reported to have an influence on SRL and root mass fraction (Freschet *et al*., 2015) and on RTD (Schaffer-Morrison *et al*., 2024). We detected no influence of soil properties, but our soil data (texture, carbon and nitrogen concentrations) mostly reflected regional and plot differences; we did not quantify fine-scale soil properties within plots. Like Weemstra *et al*. (2021), our results indicate a potential influence of fine-scale soil heterogeneity (at the centimeter level), such as soil coarse fragment load/stone content, though this was not measured here.

ITV-environment relationships are only comparable when the extent of the environmental gradient studied is the same. For instance, the influence of a given abiotic factor (e.g., water availability, temperature) on root traits and ITV can only be fully assessed when a broad gradient is considered. This can be achieved by studies spanning global or continental scales (e.g., Cardou *et al*., 2022; Han *et al*., 2025) or, in controlled conditions such as greenhouses or manipulative experiments (e.g., Jaeger *et al*., 2025). Nevertheless, we can conclude that at the regional scale of our study, and in natural systems ( in contrast to greenhouse or monoculture experiments), the magnitude and extent of ITV depends on the species as well as on the trait – which holds true at the global scale (Han *et al*., 2025) – and likely involves drivers acting at the soil centimeter, or even the rhizosphere, scale.

### 4.3. Perspectives and concluding remarks

By documenting substantial trait variation at the seedling stage – both within and across species – and highlighting pronounced within-plot ITV, our findings suggest that seedlings (conspecific or not) can adjust their phenotypes to environmental conditions at the rhizosphere scale. Although we were unable to identify specific local drivers among those studied, this high plasticity may determine the adaptive capacity of juveniles facing climate change. To distinguish the adaptive plasticity from non-adaptive or maladaptive plasticity, future studies should link trait variation to fitness metrics such as growth and survival to evaluate its consequences for regeneration success (Rowland *et al*., 2023).

Incorporating root traits and their plasticity into mechanistic models of forest succession or terrestrial biosphere models cross scales is essential to both accurately represent root contribution to ecosystem functioning and to capture water and nutrient uptake responses to environmental variations (Norby and Iversen, 2017; Warren *et al*., 2015). This integration first requires matching scales between data and model application because root ITV–environment relationships are scale- and gradient-intensity dependent, from plant and community to landscape and global levels. Accordingly, model inputs should be collected and applied at matching spatial scales (Green *et al*., 2022). For instance, forest managers using forest landscape models (e.g., LANDIS-II, Scheller *et al*., 2007) to simulate forest management scenarios should rely on root trait data collected at the community level.

A second prerequisite is to ensure appropriate reuse of trait data, which is common practice given the challenges of measuring root traits and the expansion of large trait databases (Guerrero-Ramírez *et al*., 2021; Iversen *et al*., 2021). Therefore, it is crucial to assess when ITV matters (Albert *et al*., 2011), particularly by comparing its magnitude to interspecific variation across environmental gradients (Chacón-Labella *et al*., 2023; Green *et al*., 2022; Siefert *et al*., 2015) and by identifying highly plastic traits (e.g., root system traits) and incorporating their variability into comparative studies (McCormack *et al*., 2017). The root traits examined here, linked to seedling water uptake, can be classified into three categories to help determine the conditions for their reliable integration into comparative study or models. First, traits that are easy to measure (e.g., SRL) would be best assessed in situ. Second, traits that are harder to measure but show low ITV (e.g., RBD, PAR) could be reliably sourced from species-level databases for the appropriate ontogenetic stage. Third, traits that are both difficult to measure and highly variable (e.g., MRD, R:S) may require finding proxies to enable their use in comparative functional studies.

This classification is hypothetical and its application would only be effective under certain circumstances. For instance, the effects of the trait on the studied function must be well established, which is often the case, see Freschet and Roumet, 2017, but the direction (e.g., positive or negative impacts) and magnitude of the relationship may be uncertain or species-specific. In addition, selected traits should correlate with performance metrics like growth or survival. Some studies have investigated this relationship for drought (e.g., Anderegg *et al*., 2016; Chen *et al*., 2022; Comas *et al*., 2013; Niemczyk *et al*., 2023; O’Brien *et al*., 2017), but this is not always straightforward, particularly when considering trait trade-off and plant strategies (Bergmann *et al*., 2020; Matthus *et al*., 2025).

Overall, our study contributes to a better understanding of community-level variation in tree seedling root traits along a precipitation gradient and under contrasting light conditions. It also helps fill gaps in root trait data and outlines practical directions for integrating root traits and ITV into comparative ecological studies, trait-based frameworks, and modeling efforts.

