## Supplementary material SM1 for "Intraspecific variability rivals interspecific differences in root traits of temperate tree seedlings"

*Characteristics of the fourteen studied plots: region, light conditions (type of site), humus depth, loss of ignition organic carbon, total N and C content, texture, basal area, percentage of angiosperms, total cover of the understory vegetation.*

| **Type of site** | **Plot id** | **Latitude** | **Longitude** | **Humus depth  (cm)** | **LOI Corg  (%)** | **N  (%)** | **C  (%)** | **Texture:  sand (%)** | **Texture:  silt (%)** | **Basal area  (m²/ha)** | **Angiosperms  (%)** | **Understory cover  (%)** |
| --- | --- | --- | --- | --- | --- | --- | --- | --- | --- | --- | --- | --- |
| Region: Petawawa | | | | | | | | | | | | |
| Close | PE_C_P1 | 45.977078 | -77.46342 | 3.0 | 1.0 | 0.01 | 0.33 | 97.5 | 1.3 | 37.5 | 56.1 | 53.7 |
|  | PE_C_P2 | 45.97458 | -77.437924 | 4.0 | 8.1 | 0.26 | 4.27 | 69.5 | 24.8 | 25.4 | 54.1 | 38.7 |
|  | PE_C_P3 | 45.96858 | -77.451802 | 6.0 | 2.9 | 0.06 | 1.79 | 72.5 | 21.9 | 55.6 | 39.5 | 70.0 |
| Open | PE_O_P1 | 45.981251 | -77.469517 | 0.0 | 6.4 | 0.14 | 3.54 | 54.2 | 36.4 | 0.0 | - | 93.7 |
|  | PE_O_P2 | 45.981134 | -77.462545 | 1.0 | 12.8 | 0.25 | 6.64 | 51.9 | 38.0 | 21.0 | - | 55.0 |
|  | PE_O_P3 | 45.9971822 | -77.4576969 | 7.0 | 6.3 | 0.13 | 2.89 | 62.8 | 27.7 | 35.9 | 48.7 | 75.0 |
| Region: Kenauk | | | | | | | | | | | | |
| Close | KE_C_P1 | 45.83828 | -74.869261 | 4.0 | 8.3 | 0.14 | 2.95 | 69.6 | 24.8 | 22.4 | 55.0 | 60.0 |
|  | KE_C_P2 | 45.819723 | -74.864258 | 13.0 | 4.2 | 0.10 | 1.90 | 86.3 | 10.1 | 35.3 | 100.0 | 58.7 |
| Open | KE_O_P1 | 45.819652 | -74.862668 | 0.0 | 7.6 | 0.13 | 3.37 | 58.7 | 34.4 | 0.0 | - | 82.5 |
|  | KE_O_P2 | 45.819715 | -74.868955 | 0.0 | 8.6 | 0.13 | 3.55 | 58.5 | 34.6 | 0.0 | - | 85.0 |
| Region: Duchesnay | | | | | | | | | | | | |
| Close | DU_C_P1 | 46.86094 | -71.64908 | 3.0 | 18.4 | 0.50 | 9.19 | 77.6 | 20.6 | 36.6 | 89.2 | 30.0 |
|  | DU_C_P2 | 46.52381 | -71.38702 | 4.5 | 2.4 | 0.07 | 1.35 | 75.0 | 21.5 | 26.8 | 79.0 | 73.7 |
|  | DU_C_P3 | 46.72501 | -71.50934 | 0.5 | 5.8 | 0.19 | 2.83 | 52.4 | 30.3 | 9.8 | 93.5 | 42.5 |
| Open | DU_O_P4 | 46.87683 | -71.64365 | 4.0 | 1.0 | 0.02 | 0.41 | 93.0 | 4.1 | 5.3 | 89.2 | 100.0 |
