## Supplementary material SM2 for "Intraspecific variability rivals interspecific differences in root traits of temperate tree seedlings"

*Selected traits with their abbreviation, units, definition and their relationship with plant water uptake, and associated references (not exhaustive).*

| **Traits** | **Units** | **Definition** | **Relationship with water uptake** | **References** |
| --- | --- | --- | --- | --- |
| Maximum rooting depth (MRD) | cm | “the deepest soil depth reached by the roots of an individual plant” (Tumber‐Dávila *et al.*, 2022). | Higher rooting depth allows to reach water retained in deeper soil horizons. | Freschet and Roumet, 2017 |
| Average diameter of absorptive roots (AD) | mm | Average diameter of the ten roots fragments of the 1^st^ and 2^nd^ order | Finer root diameter implies a higher radial hydraulic conductivity and a higher penetration ability. | Baca Cabrera *et al.*, 2024; Freschet and Roumet, 2017; Olmo *et al.*, 2014 |
| Proportion of absorptive roots (PAR) | % | Total length of the roots with a diameter below 0.5 mm compared to the total length of the complete root system | The higher the proportion of absorptive roots, the higher the water uptake capacity.  The amount of fine-root biomass or root length/ unit soil volume indicates the intensity of soil exploration, and the ability of a species to compete for soil nutrients. | Freschet and Roumet, 2017; Pérez-Harguindeguy *et al.*, 2013 |
| Root branching density (RBD) | n.cm^-1^ | Number of root tips by unit of length | A greater RBD allows the exploration of a larger soil volume, increasing the probability of finding wet soil patches. | Comas *et al.*, 2014; Cusack *et al.*, 2021 |
| Root tissue density (RTD) | g.cm^-^³ | Dry mass of roots by unit of root volume | Lower RTD allows to explore larger soil volume for a given biomass investment. | Cusack *et al.*, 2021 |
| Root to shoot ratio (R:S) | - | Dry mass of roots divided by the dry mass of stems and leaves/needles | Translates the trade-off between aboveground vs belowground resources acquisition. | Freschet *et al.*, 2021b |
| Specific root length (SRL) | m.g^-1^ | Ratio of total root length of the ten root fragments of the 1^st^ and 2^nd^ order to their dry mass. | Higher SRL allows a higher absorptive area for a given biomass investment. | Freschet and Roumet, 2017 |
