## Supplementary material SM3 for "Intraspecific variability rivals interspecific differences in root traits of temperate tree seedlings"

*Mean (and standard deviation) of species traits and age. SLA: Specific Leaf Area, MRD: Maximum Rooting Depth, PAR: proportion of absorptive roots, RMF: Root Mass Fraction, R:S: Root to Shoot Ratio, SRM: Specific Root Length, RTD: Root Tissue Density, AD: Average diameter.*

|  | *Abies balsamea* | *Acer rubrum* | *Acer saccharum* | *Betula alleghaniensis* | *Betula papyrifera* | *Pinus strobus* | *Quercus rubra* |
| --- | --- | --- | --- | --- | --- | --- | --- |
| Age  (years) | 7.7  (3.2) | 5.4  (2.7) | 5.9  (2.1) | 2.0  (1) | 1.8  (0.7) | 5.1  (2.4) | 3.7  (1.6) |
| SLA  (cm²/g) | 107.6  (31.6) | 281.7  (84.1) | 310.5  (105.6) | 317.4  (114.8) | 209.7  (38.9) | 114.1  (28.8) | 261.5  (82.7) |
| MRD  (cm) | 11.8  (6.8) | 13.6  (7.1) | 16.6  (9.1) | 8.2  (5.4) | 17.6  (10.2) | 11.9  (6) | 21  (8.4) |
| PAR  (%) | 62.6  (9.3) | 60.3  (9.2) | 61.7  (9.3) | 59.4  (9.2) | 48.4  (12.5) | 51.5  (11.1) | 48.1  (12.3) |
| RMF  (g/g) | 0.23 (0.07) | 0.40  (0.1) | 0.45  (0.11) | 0.28  (0.09) | 0.24  (0.09) | 0.18  (0.06) | 0.52  (0.10) |
| R:S  (-) | 0.30  (0.12) | 0.71  (0.34) | 0.88  (0.44) | 0.42  (0.22) | 0.34  (0.18) | 0.22  (0.09) | 1.19  (0.49) |
| SRL  (m/g) | 18.4  (5.0) | 48.7  (12.2) | 43.6  (14.0) | 52.5  (16.1) | 50.0  (21.4) | 15.2  (3.4) | 26.9  (10.5) |
| RTD  (g/cm³) | 0.14  (0.06) | 0.21  (0.09) | 0.24  (0.14) | 0.41  (0.05) | 0.18  (0.09) | 0.10  (0.04) | 0.25  (0.10) |
| RBD  (n/cm) | 1.41  (0.17) | 1.82  (0.26) | 1.69  (0.29) | 2.20  (0.28) | 1.87  (0.30) | 1.52  (0.27) | 1.56  (0.38) |
| AD  (mm) | 0.67  (0.16) | 0.33  (0.08) | 0.35  (0.10) | 0.38  (0.10) | 0.38  (0.13) | 0.89  (0.19) | 0.42  (0.12) |
