## Supplementary material SM4 for "Intraspecific variability rivals interspecific differences in root traits of temperate tree seedlings"

*Estimated marginal means and standard error (SE) of the traits per species.*

| **Trait** | **Species** | **Estimated Mean** | **SE** | **Compact Letter Display from Tukey test** |
| --- | --- | --- | --- | --- |
| Root to Shoot Ratio (-) | *Abies balsamea* | 0.291 | 0.080 | b |
|  | *Acer rubrum* | 0.699 | 0.080 | c |
|  | *Acer saccharum* | 0.809 | 0.081 | c |
|  | *Betula alleghaniensis* | 0.348 | 0.095 | b |
|  | *Betula papyrifera* | 0.316 | 0.115 | b |
|  | *Pinus strobus* | 0.218 | 0.085 | a |
|  | *Quercus rubra* | 1.149 | 0.084 | d |
| Specific Root Length (m.g^-1^) | *Abies balsamea* | 19.601 | 2.105 | a |
|  | *Acer rubrum* | 49.490 | 2.106 | d |
|  | *Acer saccharum* | 42.761 | 2.127 | c |
|  | *Betula alleghaniensis* | 52.490 | 2.516 | d |
|  | *Betula papyrifera* | 49.602 | 3.112 | cd |
|  | *Pinus strobus* | 15.234 | 2.245 | a |
|  | *Quercus rubra* | 27.234 | 2.208 | b |
| Maximum rooting depth (cm) | *Abies balsamea* | 11.081 | 0.110 | bc |
|  | *Acer rubrum* | 11.638 | 0.110 | bc |
|  | *Acer saccharum* | 14.799 | 0.111 | cd |
|  | *Betula alleghaniensis* | 7.422 | 0.126 | a |
|  | *Betula papyrifera* | 12.442 | 0.148 | bcd |
|  | *Pinus strobus* | 9.757 | 0.115 | ab |
|  | *Quercus rubra* | 18.401 | 0.114 | d |
| Root Tissue Density (g.cm^-^³) | *Abies balsamea* | 0.095 | 0.008 | a |
|  | *Acer rubrum* | 0.146 | 0.008 | b |
|  | *Acer saccharum* | 0.177 | 0.008 | c |
|  | *Betula alleghaniensis* | 0.086 | 0.009 | a |
|  | *Betula papyrifera* | 0.097 | 0.011 | a |
|  | *Pinus strobus* | 0.074 | 0.008 | a |
|  | *Quercus rubra* | 0.155 | 0.008 | bc |
| Proportion of absorptive roots (%) | *Abies balsamea* | 63.444 | 1.900 | c |
|  | *Acer rubrum* | 60.304 | 1.901 | c |
|  | *Acer saccharum* | 61.251 | 1.920 | c |
|  | *Betula alleghaniensis* | 58.122 | 2.247 | bc |
|  | *Betula papyrifera* | 48.811 | 2.741 | a |
|  | *Pinus strobus* | 51.838 | 2.016 | ab |
|  | *Quercus rubra* | 48.988 | 1.986 | a |
| Average diameter of absorptive roots (mm) | *Abies balsamea* | 0.674 | 0.021 | c |
|  | *Acer rubrum* | 0.331 | 0.021 | a |
|  | *Acer saccharum* | 0.352 | 0.021 | ab |
|  | *Betula alleghaniensis* | 0.379 | 0.026 | ab |
|  | *Betula papyrifera* | 0.383 | 0.034 | ab |
|  | *Pinus strobus* | 0.892 | 0.023 | d |
|  | *Quercus rubra* | 0.422 | 0.022 | b |
| Root Branching Density (tips.cm^-1^) | *Abies balsamea* | 1.377 | 0.048 | a |
|  | *Acer rubrum* | 1.820 | 0.048 | c |
|  | *Acer saccharum* | 1.682 | 0.048 | bc |
|  | *Betula alleghaniensis* | 2.166 | 0.058 | d |
|  | *Betula papyrifera* | 1.879 | 0.073 | c |
|  | *Pinus strobus* | 1.516 | 0.052 | ab |
|  | *Quercus rubra* | 1.581 | 0.051 | b |
| Specific Leaf Area (cm².g^-1^) | *Abies balsamea* | 108.880 | 19.678 | a |
|  | *Acer rubrum* | 284.673 | 19.680 | bc |
|  | *Acer saccharum* | 307.958 | 19.782 | c |
|  | *Betula alleghaniensis* | 360.259 | 20.906 | d |
|  | *Betula papyrifera* | 257.045 | 22.700 | b |
|  | *Pinus strobus* | 119.505 | 20.040 | a |
|  | *Quercus rubra* | 267.602 | 19.947 | b |

*Estimated marginal means of the traits. Means with no letter in common are significantly different according to post-hoc Tukey HSD test (α = 0.05). ABBA: Abies balsamea, PIST: Pinus strobus, ACRU: Acer rubrum, ACSA: Acer saccharum, BEAL: Betula alleghaniensis, BEPA: Betula papyrifera, QURU: Quercus rubra.*

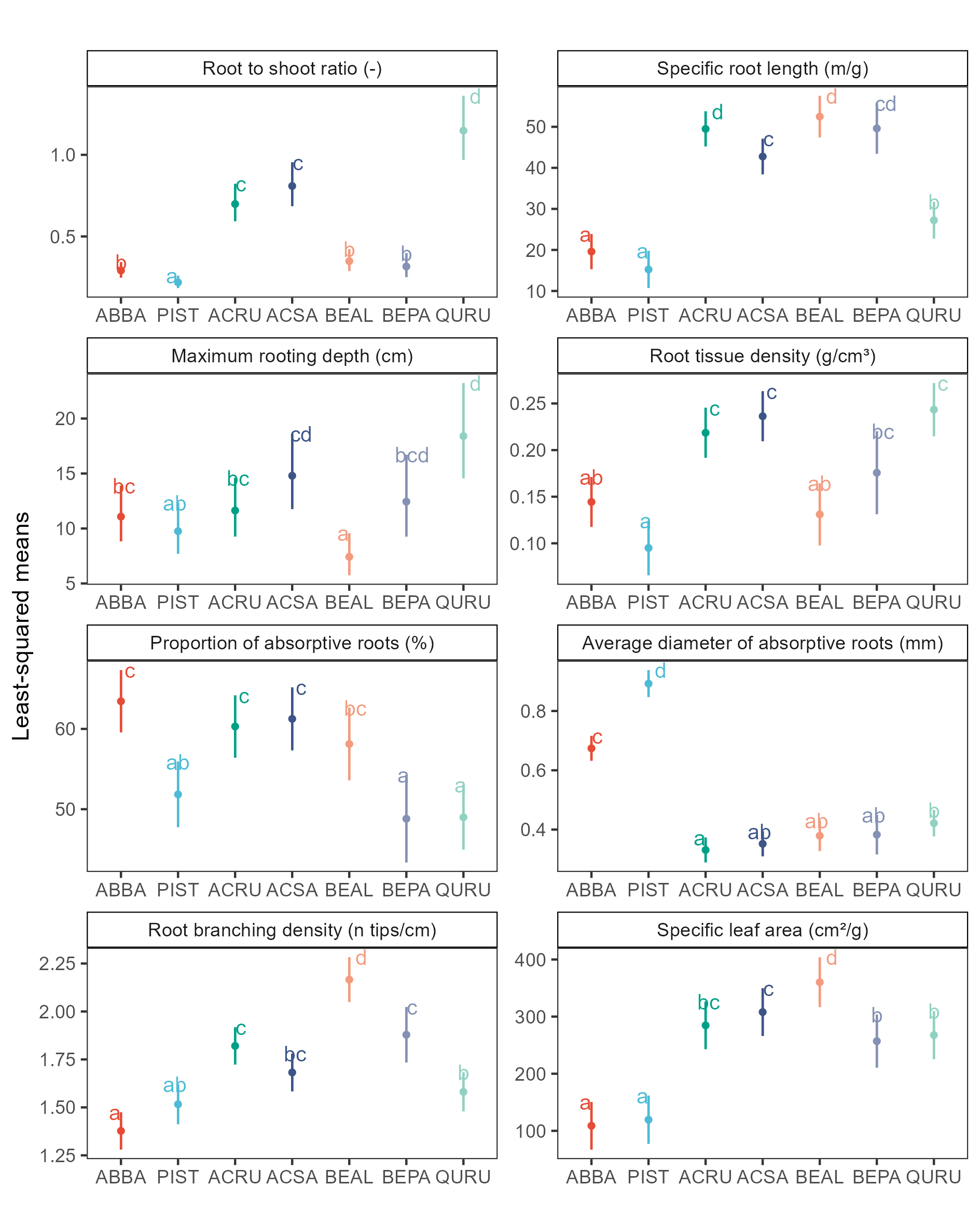
