## Supplementary material SM5 for "Intraspecific variability rivals interspecific differences in root traits of temperate tree seedlings"

*Parameter* *estimates with standard error (SE) and p-value for the fixed effects of the linear mixed models for each trait with species, plots and regions as random factors. Marginal R-squared (R²m) is the variance explained by fixed factors and Conditional R-squared (R²c) is the variance explained by both fixed and random factors. SLA: Specific Leaf Area, MRD: Maximum Rooting Depth, PAR: proportion of absorptive roots, RMF: Root Mass Fraction, R:S: Root to Shoot Ratio, SRM: Specific Root Length, RTD: Root Tissue Density, AD: Average diameter.*

| **Traits** | **Fixed effects** | **Parameter estimates** | **SE** | **p-values** |
| --- | --- | --- | --- | --- |
| **log (AD)** | Intercept (Clade: Angiosperm) | -1.02 | 0.08 | < 0.001 |
| *R²m = 0.59* | Clade: Gymnosperm | 0.75 | 0.11 | < 0.01 |
| *R²c = 0.73* |  |  |  |  |
| **log (MRD)** | Intercept | 2.46 | 0.12 | < 0.001 |
| *R²m = 0.24,* | P-PET | -0.31 | 0.05 | < 0.001 |
| *R²c = 0.48* |  |  |  |  |
| **log (PAR)** | Intercept | 3.99 | 0.06 | < 0.001 |
| *R²m = 0* |  |  |  |  |
| *R²c = 0.42* |  |  |  |  |
| **log (RBD)** | (Intercept) | 1.77 | 0.11 | < 0.001 |
| *R²m = 0.24* | Site: Open | 0.06 | 0.05 | NS |
| *R²c = 0.56* | Clade: Gymnosperm | -0.21 | 0.15 | NS |
|  | Age | -0.08 | 0.02 | < 0.001 |
|  | Site: Open x Clade: Gymnosperm | -0.16 | 0.07 | < 0.05 |
| **log (R:S)** | (Intercept) | -0.77 | 0.24 | < 0.05 |
| *R²m = 0.01* | Age | 0.07 | 0.03 | < 0.05 |
| *R²c = 0.74* |  |  |  |  |
| **RTD** | (Intercept) | 0.12 | 0.02 | < 0.001 |
| *R²m = 0.08* | Age | 0.01 | 0.00 | < 0.001 |
| *R²c = 0.68* | Clade: Gymnosperm | -0.06 | 0.02 | <0.05 |
| **SRL** | (Intercept) | 3.73 | 0.13 | < 0.001 |
| *R²m = 0.52*  *R²c = 0.77* | Site: Open | -0.09 | 0.08 | NS |
|  | Clade: Gymnosperm | -0.86 | 0.24 | < 0.05 |
|  | Age | -0.08 | 0.02 | < 0.001 |
|  | P-ETR_growing season | -0.11 | 0.04 | < 0.05 |
|  | Site: Open x Clade: Gymnosperm | -0.03 | 0.08 | NS |
| **log (SLA)** | (Intercept) | 5.93 | 0.07 | < 0.001 |
| *R²m = 0.79* | Site: Open | -0.59 | 0.07 | < 0.001 |
| *R²c = 0.87* | Clade: Gymnosperm | -1.17 | 0.10 | < 0.001 |
|  | Site: Open x Clade: Gymnosperm | 0.37 | 0.06 | < 0.001 |
