## Supplementary material SM6 for "Intraspecific variability rivals interspecific differences in root traits of temperate tree seedlings"

*Ontogeny root trait variation. Change in ranking between species from seedlings (mean per species-trait from this study) to mature trees (from TRY database; mean) for Specific Leaf Area (petiole excluded), Root Mass Fraction, Average diameter of absorptive roots, Specific Root Length and Root Tissue Density. ABBA: Abies balsamea, PIST: Pinus strobus, ACRU: Acer rubrum, ACSA: Acer saccharum, BEAL: Betula alleghaniensis, BEPA: Betula papyrifera, QURU: Quercus rubra.*

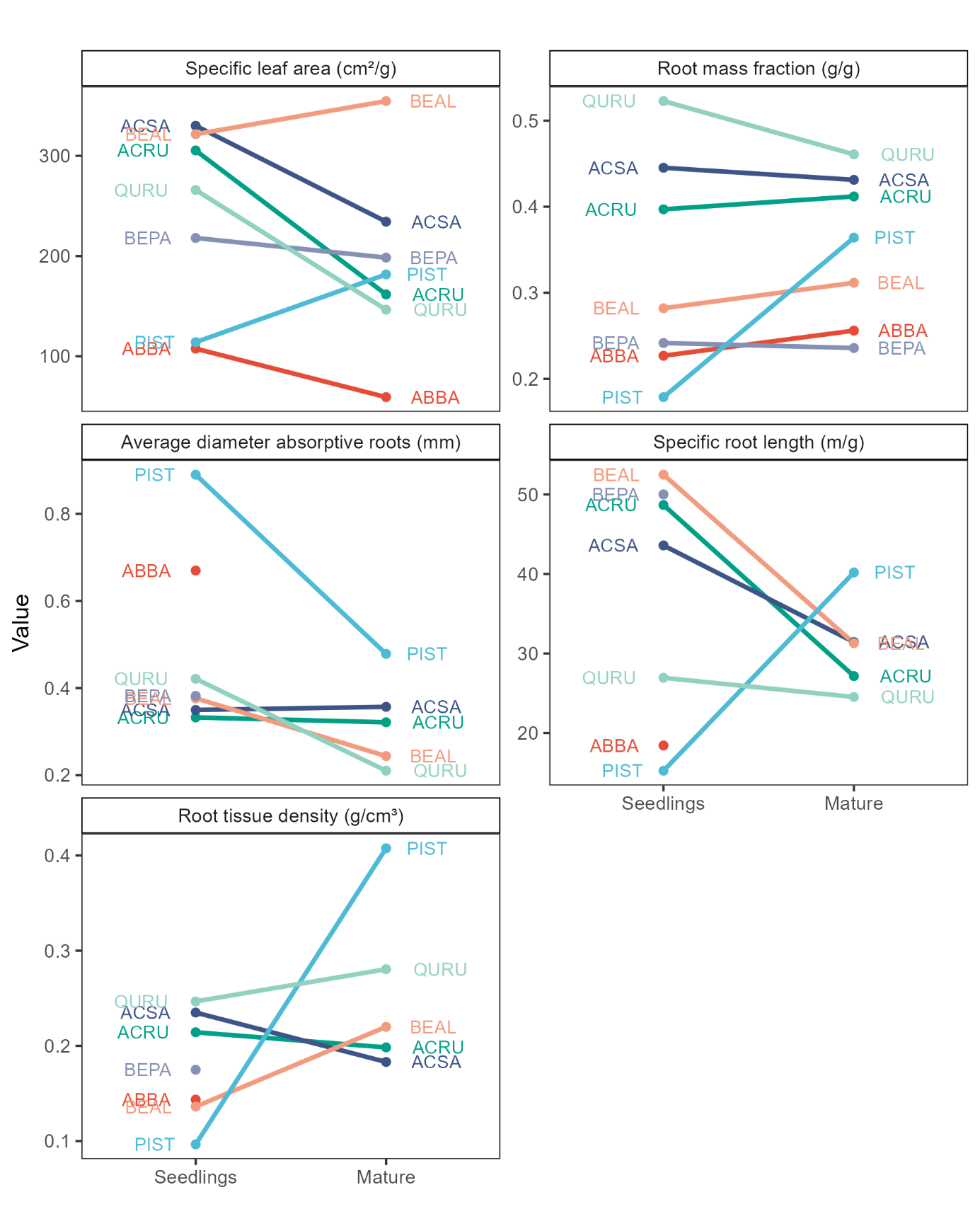
